# Desiccated cyanobacteria serve as efficient plasmid DNA carrier in space flight

**DOI:** 10.1101/2023.10.31.564425

**Authors:** Anne Kakouridis, Spencer Diamond, Thomas Eng, Heath J. Mills, Olivia Gámez Holzhaus, Michael L. Summers, Ferran Garcia-Pichel, Aindrila Mukhopadhyay

## Abstract

Effective transport of biological systems as cargo during space travel is a critical requirement to use synthetic biology and biomanufacturing in outer space. Bioproduction using microbes will drive the extent to which many human needs can be met in environments with limited resources. Vast repositories of biological parts and strains are available to meet this need, but their on-site availability requires effective transport. Here, we explore an approach that allows DNA plasmids, a ubiquitous synthetic biology part, to be safely transported to the International Space Station and back to the Kennedy Space center, without low-temperature or cryogenic stowage. Our approach relied on the cyanobacterium *Nostoc punctiforme* PC73102, that is naturally tolerant to prolonged desiccation. Desiccated *N. punctiforme* was able to carry the non-native pSCR119 plasmid as intracellular cargo safely to space and back. Upon return to the laboratory, the extracted plasmid showed no DNA damage or additional mutations and could be used as intended to transform the model Synbio host *Escherichia coli* to bestow Kanamycin resistance. This proof-of-concept study provides the foundation for a ruggedized transport host for DNA to environments where there is a need to reduce equipment and infrastructure for biological parts stowage and storage.

## Introduction

Recent advances in synthetic biology have led to the development of DNA-based systems for information storage and biomanufacturing (Rutten *et al*., 2018), holding great promise for space applications. DNA is a stable and efficient molecule for storing genetic information and is particularly attractive for space-based biomanufacturing, where products can be synthesized when and where they are needed (Panda *et al*., 2018). However, the space environment poses significant challenges to the stability and functionality of DNA-based systems because exposure to cosmic radiation, microgravity, and extreme temperatures can cause DNA damage, strand breaks, and other genetic changes (Mileikowsky *et al*., 2000; Horneck *et al*., 2001; Horneck, 2003; Moreno-Villanueva *et al*., 2017). In addition, space environmental factors can also affect the gene expression and protein synthesis machinery of cells, impacting the functionality and efficiency of DNA-based systems (Aniveka *et al*., 1990; de la Torre *et al*., 2010). Therefore, to realize the full potential of DNA-based biomanufacturing in space, it is essential to gain a comprehensive understanding of the effects of space environmental factors on DNA stability and function. Advancing our knowledge in this area will enable the development of advanced DNA-based technologies for space applications, including the production of materials, medicines, and other biomolecules required for sustaining human exploration and habitation in space. In this study, we address the need for robust methods to safely transport DNA to space stations.

To protect DNA from the stresses associated with spaceflight while safely carrying it at room temperature, we experimented with the use of cyanobacteria, well-known for native desiccation tolerance (Inoue-Sakamoto *et al*., 2018), as DNA carriers. *Nostoc punctiforme* is a model cyanobacterial system that is sufficiently genetically tractable to examine its potential to serve as transport chassis for DNA (Thiel & Poo, 1989). Moreover, as a fully autotrophic organism, it enables a simpler feedstock logistical supply chain requiring only sunlight and CO_2_ as inputs, where the CO_2_ will always be generated as a consequence of human activity in these space environments (Averesch *et al*., 2023). In this study, we probed *N. punctiforme*’s capacity to act as a rugged DNA carrier in a real space flight test involving travel as cargo to the International Space station, subsequent month-long orbit, and eventual return to the laboratory to be examined for revival, analyses of DNA integrity and rate of mutations (**Figure 1**).

**Figure 1:**
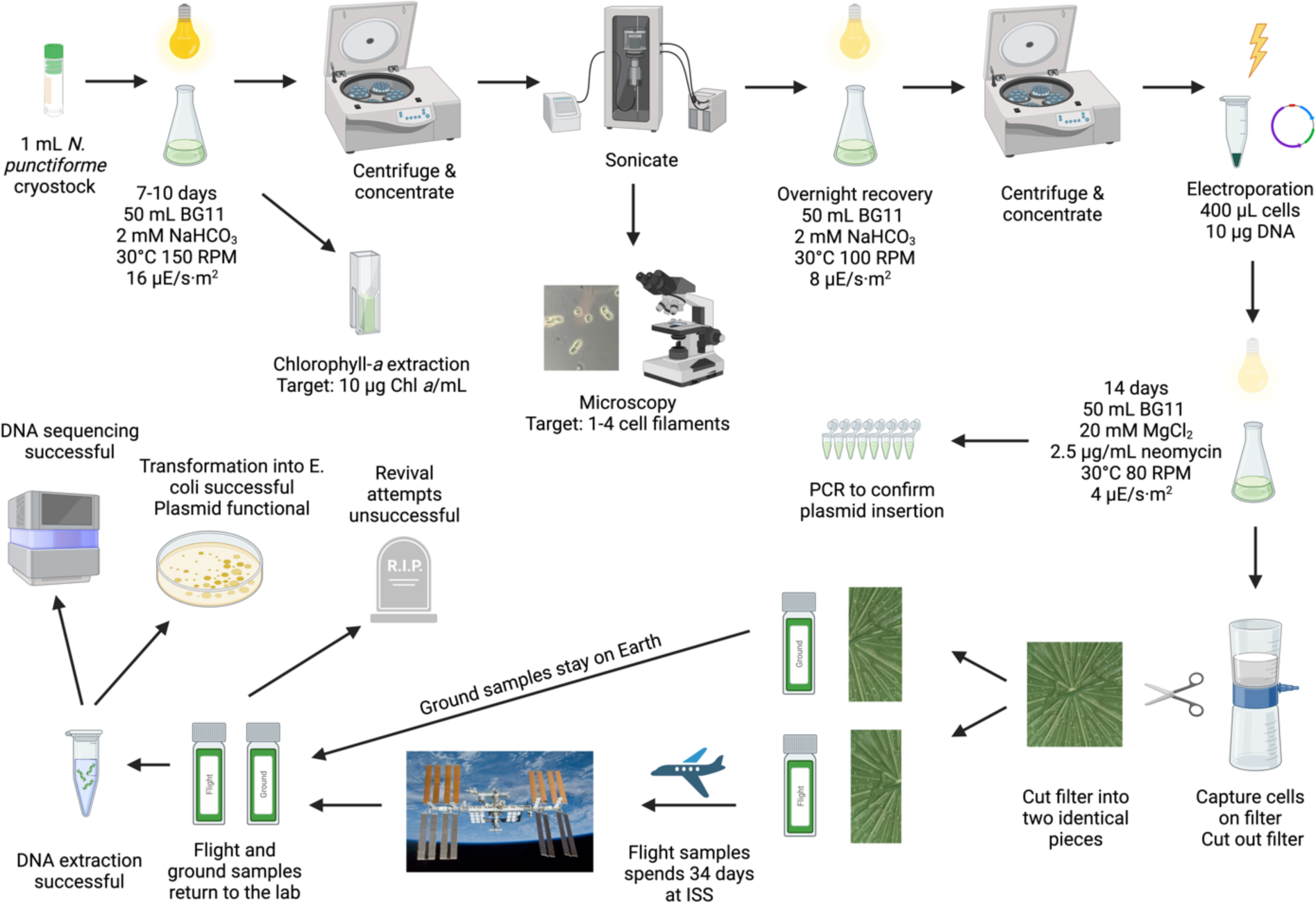
Workflow

## Materials and Methods

### Growing conditions

*N. punctiforme* (ATCC 29133/PCC 73102) (**Figure 2a**) was obtained as a cryostock (5% DMSO final concentration) as routinely kept in the Garcia-Pichel laboratory at Arizona State University, Tempe, AZ, USA. Each 1 mL cryostock was thawed and transferred to a 250 mL baffled flask containing 50 mL BG11 minimal medium containing nitrate. The 1X BG11 medium used in this study was made by diluting 50X Cyanobacteria BG-11 Freshwater solution (Cat. No. C3061, Sigma-Aldrich, Inc, St. Louis, MO, USA) with autoclaved milliQ water. Flasks were placed on a shaker at 150 RPM, 30°C, under 24-hour lights emitting 16 μE/s·m^2^ (Mosthink LED Plant Grow Lights Strips Full Spectrum, Amazon, Seattle, WA, USA). The light intensity the cultures experienced was checked weekly by measuring photosynthetically active radiation (PAR) with a PAR meter (PHOTOBIO Advanced Quantum PAR Meter by Phantom, Amazon, Seattle, WA, USA). To test different growing media amendments, some *N. punctiforme* cryostocks were thawed and grown in BG11 supplemented with NaHCO_3_ (final concentrations 1 mM, 2 mM, 4 mM, 6 mM, 8 mM, and 10 mM) (**Table S-1**).

**Figure 2:**
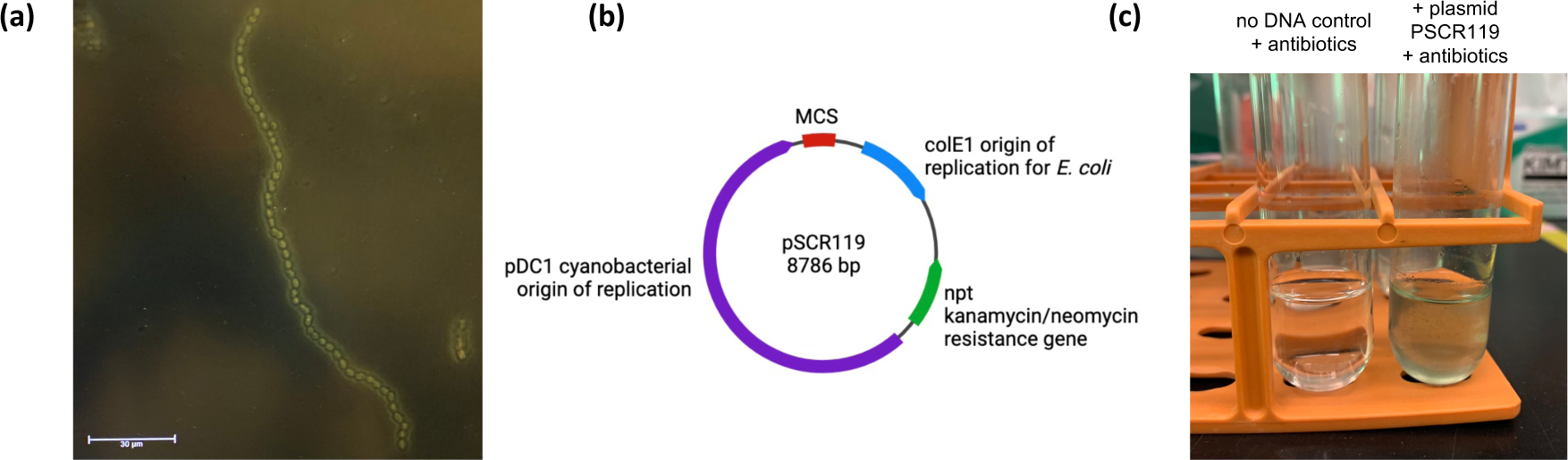
**(a)** Microscopy image of WT *N. punctiforme* **(b)** Plasmid map of pSCR119 **(c)** *Nostoc punctiforme* growing in BG11 supplemented with kanamycin (2.5 µg/mL final concentration) 10 days after transformation when pSCR229 was not added (left) vs added (right).

Dense cultures were obtained after 7-10 days of growth. Cell density was measured by Chlorophyll-*a* (Chl *a*) extraction following a protocol from the Summers laboratory at California State University, Northridge, CA, USA. In brief, 1 mL of each culture was transferred to a microcentrifuge tube and centrifuged at 15,000 RPM at room temperature for 1 min. Then, 900 µL of supernatant were removed from each tube and replaced with 900 µL of 100% methanol. Each tube was vortexed to break up the pellet, allowed to sit for 5 min at room temperature in the dark, then was vortexed again. Tubes were again centrifuged at 15,000 RPM at room temperature for 1 min. The supernatant of each tube was transferred to a cuvette and absorbance was read at 665 nm against a 90% methanol blank. The absorbance value was multiplied by 12.7 to obtain µg Chl *a*/mL (based on the extinction coefficient, Meeks & Castenholz, 1971) and cell density (1 µg/mL Chl *a* = ∼ 10^6^ cells). Our target cell density was 10 μg Chl *a*/mL for a 50 mL culture to prepare electrocompetent cells.

### Transformation conditions

Plasmid pSCR119 (Summers *et al*., 1995) (**Figure 2b**) was obtained from stocks kept in the Summers laboratory. This shuttle plasmid is compatible with both *E. coli* and *N. punctiforme* and confers kanamycin/neomycin (kan/neo) resistance. The plasmid DNA was transformed into *E. coli* following a standard heat shock protocol then isolated by miniprep (Cat. No. 27104, QIAprep Spin Miniprep Kit, Qiagen, Germantown, MD, USA) following the manufacturer’s instructions and eluted in molecular grade water to minimize the salt concentration for electroporation into *N. punctiforme*.

Transformations by electroporation were performed following a protocol by Summers *et al*. (1995) with some modifications (**Table S-2**). In brief, each 50 mL culture was transferred to a 50 mL falcon tube and centrifuged at 4000 RPM for 5 min. The supernatant was removed so that each culture was concentrated down to 5 mL. The 5 mL concentrated cultures were sonicated to break up the *N. punctiforme* filaments using a 2 mm microtip, amplitude 1, and the following settings were tested: 10 × 10 sec bursts, 5 × 10 sec bursts, 4 × 5 sec bursts or 3 × 5 sec bursts (**Table S-2**). After sonication, a droplet of each culture was mounted on a glass slide and observed under a compound microscope to confirm that most *N. punctiforme* filaments were down to 1-4 cells in length. The 5 mL concentrated cultures were individually transferred to 45 mL BG11 in 250 mL flasks and allowed to recover overnight at 50% light intensity, 100 RPM, 30°C. The next morning, the recovered cultures were centrifuged at 4000 RPM for 5 min then supernatant was removed. The pellets were washed with 40 mL autoclaved room temperature milliQ water 4 times, centrifuging each time. After the fourth wash, the pellets were resuspended to 5 mL in room temperature milliQ water. In microcentrifuge tubes, 400 µL of concentrated washed cells and 10 µg of plasmid DNA were mixed by pipetting and allowed to incubate on ice for 1 min. The mixture was transferred to a 2 mm cuvette for electroporation with the following parameters: 600 Ω, 1.6 kEV, 25 μF. The goal was to obtain a time constant of 11.5-13.5 ms. Immediately after electroporation, 1 mL of BG11 supplemented with MgCl_2_ (20 mM final concentration) was added to the microcentrifuge tube, then the content of the tube was transferred to 50 mL of the same media in a 250 mL baffled flask. All the flasks were allowed to recover for 24 hours at 25% light intensity, 80 RPM, 30°C. Cultures were then centrifuged again and transferred to fresh BG11 with either kanamycin or neomycin was added to the flasks to reach a final concentration of 1.25 or 2.5 or 5 or 10 µg/mL (different concentration were tested) and the flasks were incubated at 25% light intensity, 80 RPM, 30°C for 14 days.

To check for successful transformation, the cell density of transformants and control (no plasmid added) was measured every few days by Chl *a* extraction as described previously to confirm that transformants were growing more than wild type (WT) cultures in the presence of kan/neo (**Figure 2c**). In addition, the presence of the kan/neo resistance genes in transformant cultures was checked every few days by PCR, gel electrophoresis, and Sanger sequencing. To do so, 1 mL of each culture was pelleted by centrifugation at 8000 rpm for 3 min, then the supernatant was removed. The pellets were resuspended in 50 µL BG11 and 1 µL was then used as the DNA template for PCR. PCR was conducted following the manufacturer’s instructions for the Q5® High-Fidelity 2X Master Mix (Cat. No. M0492S, New England BioLabs, Inc, Ipswich, MA, USA), the following conditions: 98°C for 30 sec and [98 °C for 10 sec, 71 °C for 5 sec, 72 °C for 30 sec] × 35 cycles, and the following primers: forward 5’-CTGCAATGATACCGCGAGACCC-3’, reverse 5’-CCAGTCCGCAGAAACGGTGC-3’ with an expected PCR product size of 1288 bp for gene encoding the kan/neo resistance marker gene.

### Sample preparation for transportation to the International Space Station

Successful transformants and WT *N. punctiforme* cultures were sterile filtered onto 0.22 µm cellulose acetate membranes in Corning® 150 mL Bottle Top Filter (Cat. No. 430626, Corning Inc, Corning, NY USA). The filters were cut out of the bottle top using autoclaved metal scissors and tweezers and placed into petri dishes, then allowed to dry overnight in the dark at room temperature (**Figure 3a**). The following day, each filter was cut in half, and each half-filter was placed in a Rhodium Cyrotube 0005 (NASA Part #: RhCT-0005) (**Figure 3b**). These Rhodium proprietary tubes provide 4 mL working volume for samples and have been NASA flight safety certified for operation on the ISS. Filters were loaded into the RhCT-0005s using autoclaved metal scissors and tweezers.

**Figure 3:**
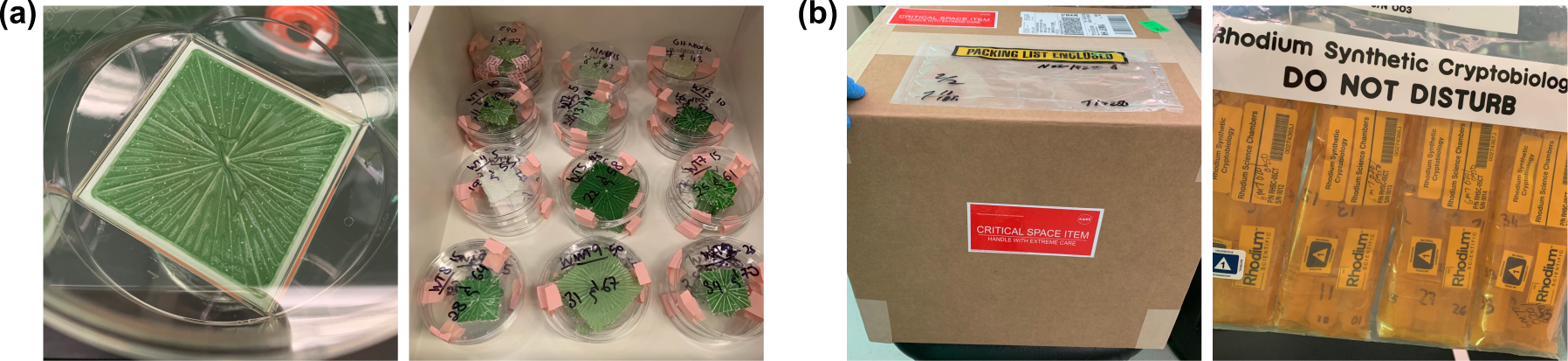
**(a)** Images of filters with a layer of wild type and transformed *Nostoc punctiforme* **(b)** Image of cells in cryo-tubes packaged for shipment to and from the Kennedy Space Center and the International Space Station.

Since the limitation of cell quantity may be different for resuscitation and DNA extraction after a time period as cargo in during space travel, we prepared several concentrations of cell mass on filter paper from both transformed and WT *N. punctiforme* (**Table 1**). With this strategy, we produced filters carrying 3,302,000 to 14,986,000 cells for transformed *N. punctiforme* and 1,240,000 to 445,770,000 for WT *N. punctiforme* (**Figure 3**, **Table 1**).

**Table 1:** Flight and Ground sample list. Corresponding Flight and Ground samples (e.g., 1 & 37) are two halves of the same filter. Wild type samples were used to test different DNA extraction and revival methods upon samples returning to the laboratory.

| Ground sample | Flight sample | Description | # cells/mL | Volume filtered (mL) | Total # cells on filter | Transform into *E. coli* |
| --- | --- | --- | --- | --- | --- | --- |
| 1 | 37 | Transformant (pSCR119) | 419100 | 20 | 8,382,000 | Yes |
| 2 | 38 |  | 736600 | 20 | 14,732,000 | Yes |
| 3 | 39 |  | 825500 | 5 | 4,127,500 | Yes |
| 4 | 40 |  | 825500 | 15 | 12,382,500 | Yes |
| 5 | 41 |  | 1231900 | 2.5 | 3,079,750 | No |
| 6 | 42 |  | 1231900 | 7.5 | 9,239,250 | Yes |
| 7 | 43 |  | 660400 | 5 | 3,302,000 | Yes |
| 8 | 44 |  | 660400 | 12.5 | 8,255,000 | Yes |
| 10 | 46 | Wild type | 3162300 | 5 | 15,811,500 |  |
| 11 | 47 |  | 3162300 | 5 | 15,811,500 |  |
| 12 | 48 |  | 3162300 | 5 | 15,811,500 |  |
| 13 | 49 |  | 3403600 | 2.5 | 8,509,000 |  |
| 14 | 50 |  | 3403600 | 2.5 | 8,509,000 |  |
| 15 | 51 |  | 3403600 | 2.5 | 8,509,000 |  |
| 16 | 52 |  | 7835900 | 5 | 39,179,500 |  |
| 17 | 53 |  | 7835900 | 5 | 39,179,500 |  |
| 18 | 54 |  | 7835900 | 5 | 39,179,500 |  |
| 19 | 55 |  | 496000 | 2.5 | 1,240,000 |  |
| 20 | 56 |  | 496000 | 2.5 | 1,240,000 |  |
| 21 | 57 |  | 496000 | 2.5 | 1,240,000 |  |
| 22 | 58 |  | 6019800 | 25 | 150,495,000 |  |
| 23 | 59 |  | 4038600 | 12.5 | 50,482,500 |  |
| 24 | 60 |  | 4038600 | 12.5 | 50,482,500 |  |
| 25 | 61 |  | 4622800 | 7.5 | 34,671,000 |  |
| 26 | 62 |  | 4622800 | 7.5 | 34,671,000 |  |
| 27 | 63 |  | 4622800 | 7.5 | 34,671,000 |  |
| 28 | 64 |  | 2959100 | 7.5 | 22,193,250 |  |
| 29 | 65 |  | 2959100 | 7.5 | 22,193,250 |  |
| 30 | 66 |  | 2959100 | 7.5 | 22,193,250 |  |
| 31 | 67 |  | 1498600 | 25 | 37,465,000 |  |
| 32 | 68 |  | 1498600 | 25 | 37,465,000 |  |
| 33 | 69 |  | 1498600 | 10 | 14,986,000 |  |
| 34 | 70 |  | 1955800 | 12.5 | 24,447,500 |  |
| 35 | 71 |  | 1955800 | 12.5 | 24,447,500 |  |
| 36 | 72 |  | 2971800 | 150 | 445,770,000 |  |

All samples were air-couriered from Lawrence Berkeley National Laboratory (LBNL), Berkeley, CA, to the Kennedy Space Center (KSC), Merritt Island, FL, USA on December 14, 2021. Upon arrival, Rhodium Scientific packed the RhCT-0005s containing samples into two sets of four Rhodium Science Chamber 05CT (NASA Part #: RhSC-05CT) facilities. Each facility contained 9 samples contained within two layers of Bitran^(R)^ bags, labeled following NASA approved labeling plan, then grouped in sets of 4 into gallon-size plastic storage bags. This configuration was flight safety certified by Rhodium for integration and operation on the ISS. One set of samples, hereafter referred to as the ‘Ground’ samples, remained at KSC, while the other, identical, set of samples, hereafter referred to as the ‘Flight’ samples, launched to the International Space Station (ISS) on the SpaceX Crew Dragon Commercial Resupply Service Mission 24 (CRS-24) on December 21, 2021. Rhodium monitored astronaut on orbit stowage and retrieval activities from Rhodium Mission Control in Houston; a facility with video and audio links routed from the ISS through NASA Marshall Space Flight Center. The flight samples remained on the ISS undisturbed within the Japanese Experiment Module for 34 days before splashdown and return to KSC on January 24, 2022. Flight and Ground samples were then shipped to LBNL.

### Sample rehydration, DNA extraction and sequencing

Flight and Ground transformant and WT samples were taken out of the tubes and placed in petri dishes using autoclaved tools. To attempt reviving the samples, 2 mL of BG11 were pipetted onto the filters and the samples were incubated overnight at room temperature in the dark. The cells were then gently scraped off the filter and resuspended in BG11 using sterile inoculation loops. The BG11 containing resuspended cells were transferred to a glass tube and 3 mL of additional BG11 was added. The tubes were placed at 25% light, 80 RPM, 30°C. Another way the samples were attempted to be revived was by placing filters directly into tubes containing 5 mL of BG11 and placed on a shaker at 25% light, 80 RPM, 30°C.

To directly extract DNA from the desiccated Flight and Ground transformants and WT samples, we experimented with different methods. First, we placed filters in 2 mL tubes with beads and 1 mL TE buffer (Qiagen plasmid isolation kit, Qiagen Germantown, MD, USA) and ran them in the tissue lyser at either ½ speed or max speed for 30 sec or 1 min. Then, we followed a standard phenol:chloroform DNA extraction (Griffiths *et al*., 2000; Sambrook & Russel, 2006). Another way we attempted to extract DNA from the desiccated samples was by rehydrating filters in BG11 for 15 min, 2 hours, 12 hours, 24 hours and 48 hours, then gently scraping the cells of the filters with inoculation loops and placing the resuspended cells into the first tube of the DNeasy PowerSoil Pro Kit (Cat. No. 47016, Qiagen, Germantown, MD, USA) containing beads then following the manufacturer’s instructions. We also placed filters whole or cut into different size pieces (2 × 2 mm, 4 × 4 mm, 6 × 6 mm, 8 × 8 mm) in the first tube of the DNeasy PowerSoil Pro Kit using autoclaved scissors and tweezers. 500 µL of buffer (10 mM Tris-Cl. 0.5 mM EDTA; pH 8.0) was added and samples were allowed to incubate for 15 min, 2 hours, 12 hours, 24 hours and 48 hours in the dark at room temperature. Then, samples were vortexed at different speed and length of time: 10 min at full speed, 10 min at half speed, 15 min at half speed. The rest of the protocol followed the manufacturer’s instructions.

The quantity and quality of the extracted raw DNA were measured by nanodrop spectrophotometer (Thermo Fisher, USA) per manufacturer’s instructions. 100 ng of DNA was transformed into NEB^®^ 10-beta Competent *E. coli* (High Efficiency) (Cat. No. C3019H, New England BioLabs, Inc, Ipswich, MA, USA) following the manufacturer’s instructions with a recovery time of either 1 hour (standard time) or increased to 2 or 3 hours. For each sample, the entire reaction was plated onto selective LB plates. Plates were incubated overnight at 37°C (**Figure S-1**).

For each plate, colonies were counted (**Table S-3**) then scraped off the plate and pooled in 1 mL LB Kan media. The pooled colonies were resuspended and mixed by pipetting, then 500 µL were used for DNA isolation by miniprep (Cat. No. 27104, QIAprep Spin Miniprep Kit, Qiagen, Germantown, MD, USA). The extracted DNA was quantified using the Qubit™ dsDNA BR Assay Kit (Cat. No. Q32850, ThermoFisher Scientific, Waltham, MA, USA) and its purity was determined with a NanoDrop spectrophotometer (**Table S-4**). Raw plasmid DNA and isolated plasmid DNA were sent for Illumina sequencing at QB3 Genomics (UC Berkeley, Berkeley, CA) (**Table S-4**).

### DNA sequence analysis

Illumina adapter sequences and phiX sequence were removed from raw reads using BBduk (https://sourceforge.net/projects/bbmap/) with default parameters. Reads were subsequently quality filtered using Sickle (https://github.com/najoshi/sickle) with default parameters. Trimmed and quality filtered reads from each sample (mean depth ∼ 98M 150bp reads per sample) were mapped back to a combined reference sequence file containing the *N. punctiforme* genome (Taxonomy ID: 63737) and the pSCR119 plasmid (Taxonomy ID: 282192) using bowtie2 with default parameters (Langmead & Salzberg, 2012). Mapped read files from bowtie2 were converted into sorted indexed bam format using samtools (Danecek *et al*., 2021). Reference sequence coverage and divergent nucleotide sites were quantified from bam files using inStrain profile within the inStrain package (Olm *et al*., 2021). Divergent nucleotide sites are the sum of single nucleotide substitutions (SNS), a fixed nucleotide change supported by all reads relative to the reference sequence, and single nucleotide variants (SNV), a nucleotide change relative to the reference sequence supported by a significant sub-fraction of the mapped reads. InStrain as was run with default parameters which required ≥ 95% alignment identity to a reference for a read to be used in the analysis, the minimum coverage to call a variant to be ≥ 5X, and the minimum fraction of reads with a variant to call an SNV to be ≥ 5%.

Outputs from inStrain were combined and analyzed using a custom R script (**Supplementary Data S-1**). We first assessed the coverage and divergent site count estimated by InStrain for the *N. punctiforme* genome and pSCR119 plasmid in each sample and each pair of samples retained on the ground (ground) or sent to space (flight). We observed very low coverage, likely due to inefficient genomic DNA extraction from lyophilized samples, for the *N. punctiforme* genome and thus limited our analysis to divergent sites in the pSCR119 plasmid sequence. For analysis of divergent sites in the pSCR119 plasmid sequence we only retained samples for analysis that had a complete ground-flight pair and where the difference in coverage between samples in a pair was ≤ 10-fold (**Figure S-5**). All plotting was performed using ggplot2 in R (Wickham, 2011), and statistical testing to evaluate significant differences in divergent sites between ground and flight sample pairs was performed in R using both the student’s t-test and the Wilcoxon rank sum test.

## Results

In this study, we used the innate nature of cyanobacteria for desiccation and resuscitation to provide a potential rugged transport workflow where no refrigeration or freezing is required. *N. punctiforme* provided an ideal model system because it has published cultivation and transformation methods. However, prior to conducting DNA transformation or desiccation workflows, it was necessary to optimize strain cultivation and maintenance in the laboratory using reported conditions. While many different reports exist to cultivate *N. punctiforme* (e.g., Meeks, 1998; Nowruzi *et al*., 2013; Guljamow *et al*., 2017), the condition that worked best in our hands was supplementing BG11 media with NaHCO_3_ (final concentration 2 mM) to produce the most robust biomass. Results from supplementing BG11 with different concentrations of NaHCO_3_ are provided in **Table S-1**.

Next, we optimized the conditions that allow best transformation outcomes. To test this, we explored published protocols (e.g., Summers *et al*., 1995) with some variations described in the Methods section. We found that among the conditions tested reported in the Methods section and **Table S-2**, the following conditions produced the greatest transformation efficiency: 3×5 sec bursts sonication, 10 µg plasmid DNA, and either 2.5 or 5 µg/mL kan or neo. We found that maximal growth was impacted by transformation, and conditions required for maintenance of plasmid DNA. Here also we tested several regimes of antibiotic amendments to obtain the ideal tradeoff between cell mass and retention of plasmid DNA (**Table S-2**).

Rehydration and DNA extraction after a period of desiccation was the key aspect of discovery for this study. *A priori* it was not known what length of time could be accommodated before sample revival was no longer observed. Further the quantity of cells required for successful rehydration were also unknown, thus we explored desiccation rehydration workflows using both transformed and WT *N. punctiforme*. All approaches attempted are described in detail in the Methods section. In our pre-flight tests, we were able to revive both transformant and WT cells after up to four weeks of desiccation on filters. However, we were unfortunately not able to revive transformants or WT desiccated cells from Flight or Ground samples using any of the protocols we attempted.

We were successful in extracting DNA from transformant and WT desiccated cells from Flight and Ground samples. Among the different methods we tested as detailed in the Methods section, we achieved greatest DNA yield and quality using the following steps: (1) cut the filters into 2 × 2 mm size pieces, (2) place the pieces in the first tube of the DNeasy PowerSoil Pro Kit with 500 µL of buffer (10 mM Tris-Cl. 0.5 mM EDTA; pH 8.0), (3) incubate the tubes for 12 hours in the dark at room temperature, (3) vortex the tubes for 15 min at half speed, and (5) follow the rest of the kit manufacturer’s instructions. Other conditions (e.g., larger pieces of filter, shorter incubation times in buffer, different vortexing regimes) resulted in lower DNA yield and quality. It is worth noting that > 3,000,000 cells on a 4-cm^2^ piece of filter seemed to be the lower limit for successful DNA extraction regardless of the method used.

A portion of the plasmid DNA extracted from the desiccated cell was sent for sequencing. Another portion of the extracted DNA was used for transformation into *E. coli*. The transformed *E. coli* cells were able to grow on media containing kan/neo (final concentration 50 µg/mL), indicating that the plasmid had retained its functions in terms of replication potential and conferring kan/neo resistance. We then pooled all the colonies per plates and extracted the plasmids from *E. coli* to obtain more DNA. This DNA was also sent for sequencing.

Sequencing of both the plasmid DNA directly extracted from the desiccated cells and the plasmid transformed and extracted from *E. coli* revealed no significant differences in the numbers of SNPs for Flight and Ground samples (**Figure 4, Figures S-2, S-3, and S-4**).

**Figure 4:**
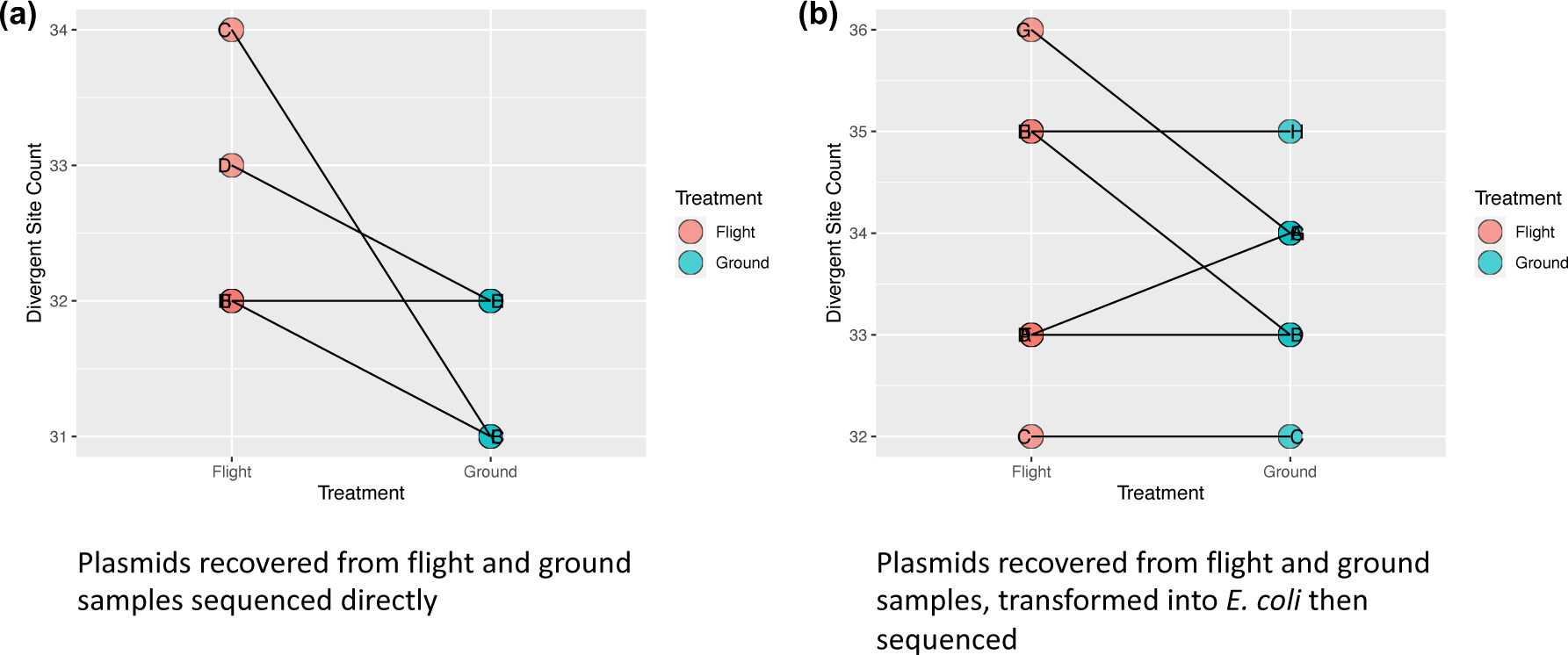
Comparison of divergent site counts for plasmids extracted from *Nostoc punctiforme* from 8 Flight and 8 Ground samples and **(a)** sequenced directly and **(b)** transformed into *E. coli* then extracted and sequenced. **(a,b)** Sequences from Flight and Ground samples were compared for SNPs. Some of the samples (dots) overlap in the figure due to having the same number of SNPs.

## Discussion

The main goal of this study was to establish a model cyanobacteria such as *N. punctiforme* as an efficient DNA transport system for travel where cargo capacity and infrastructure needs to be minimal. One such configuration is space travel, where the use of large cooling equipment can be both capacity and cost prohibitive. The cyanobacterial strain, *N. punctiforme* was chosen for this endeavor due to its natural tolerance to desiccation. Known to be the primary components in biological soil crusts (BSCs; Belnap, 2001; Belnap *et al*., 2003) in arid and semiarid regions worldwide (Weber *et al*., 2022), a unique feature of BSCs is the ability of the constituent microorganisms, particularly cyanobacteria, to withstand extended periods of drought and to revive rapidly upon rehydration (Rajeev *et al*., 2013; Garcia-Pichel, 2023). Such cyanobacteria have evolved various physiological and morphological adaptations that enable them to survive desiccation and persist for long periods in a dormant state (Dabravolski *et al*., 2013) and rapidly resume metabolic activity and photosynthesis when hydrated (Garcia-Pichel, 2023). In this context, we examined the transported and desiccated Nostoc strains for resuscitation and growth. In our workflow, the desiccated *N. punctiforme* that remained in desiccated state for 34 days at the ISS and 60 days in total, were not able to grow upon hydration and cultivation in the laboratory. Since the ground control samples for these also could not be resuscitated, we concluded that the length of time was too long with the methods employed. However, our limited samples only allowed a subset of conditions to be tested to attempt cell resuscitation. In the future, when more such opportunities are available (this study was limited to one journey to the ISS), the use of different desiccation parameters (e.g., via slower desiccation or pelleting vs filtration) and resuscitation approaches (e.g., in a wider range of cultivation media) could provide methods that revive the carrier Nostoc strains.

The extracted DNA showed negligible SNPs suggesting that the lack of revival of the strain itself may have other epigenetic causes. More importantly, the fidelity of the plasmid DNA was conserved with few to no mutations and consistent with our ability to transform into a model synbio host *E. coli* and demonstrate function. We conclude that while *N. punctiforme* cultures were not able to be revived after space travel, they were able to protect the plasmid DNA from any deleterious mutations during the space travel conditions they experienced while being transported at room temperature.

This study represents a key step towards a much larger set of goals being envisioned for microbiology enabled synthetic biology in space (Horneck *et al*., 2010). In this study we explored the basic but essential molecular biology workflow; that of DNA parts transport and availability. The ability to transport a plasmid DNA without mutations, without the need for refrigeration or cryo-storage and successfully transform a common synthetic biology workhorse after exposure to space conditions sets the stage for many synthetic biology workflows. The use of *N. punctiforme* was important and showed that microbes like these can remain desiccated for extended periods in the presence of radiation and permit minimal damage to the harbored plasmid DNA. This study provides an immediately usable workflow for non-native DNA transport using desiccated Nostoc strains and future applications that use the Nostoc itself in space synthetic biology applications.

## Data Availability

Data is submitted to NCBI SRA (SubmissionID: SUB13920154 BioProject ID: PRJNA1034103) and also available upon request.

## Acknowledgments

We thank Kevin Klicki for helping us obtain the *Nostoc* strains and plasmids, Alex Codik for archiving strains and plasmids, and Juliana Artier and Andreja Kust for their thoughtful advice on cyanobacteria culturing. We also thank Dr. RP Oats for the initial work in drafting the proposal for this study and Lucas Waldburger for help with uploading data to public repositories.

