## Supplementary Information for "Desiccated cyanobacteria serve as efficient plasmid DNA carrier in space flight"

**Figure S-1**: Images of plates after transforming raw DNA extracted from Ground and Flight samples into *E. coli* as described in the Methods section. Numbers correspond to samples indicated in Table 1.

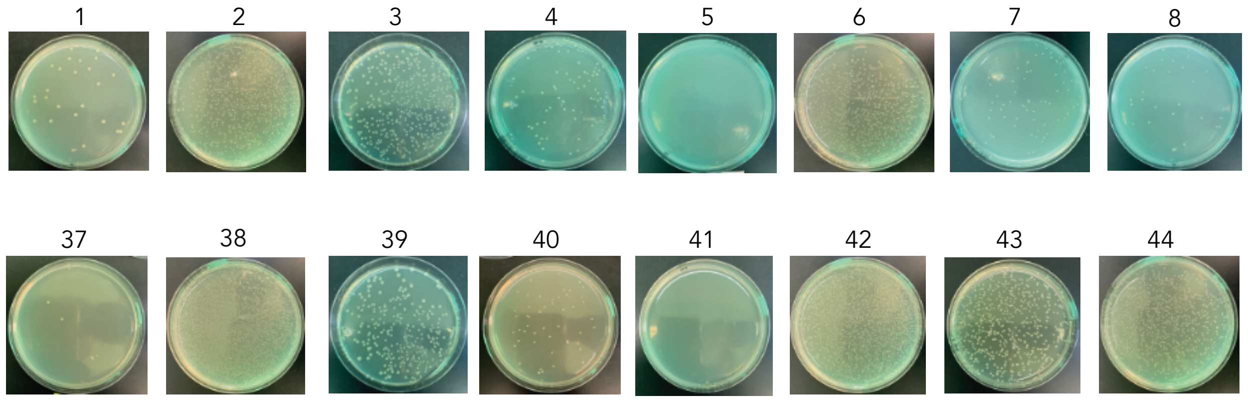

Figure **S-2**: Assessment of coverage for plasmid pSCR119 directly extracted from Flight and Ground samples and sequenced.

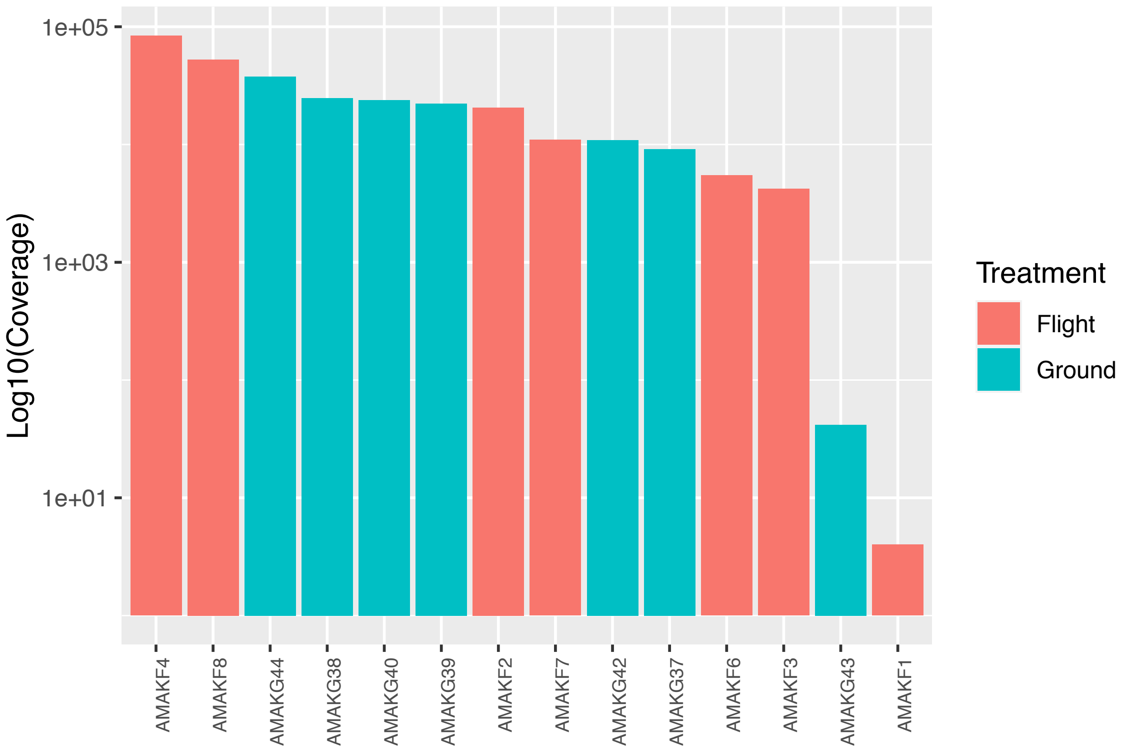

Figure **S-3**: Assessment of coverage for plasmid pSCR119 directly extracted from *N. punctiforme* flight and ground samples and sequenced. In general coverage of the plasmid in DNA extracts of *N. punctiforme* was good. There were only 2 sample pairs (A and G) where the depth of sequencing coverage was highly divergent between the pair.

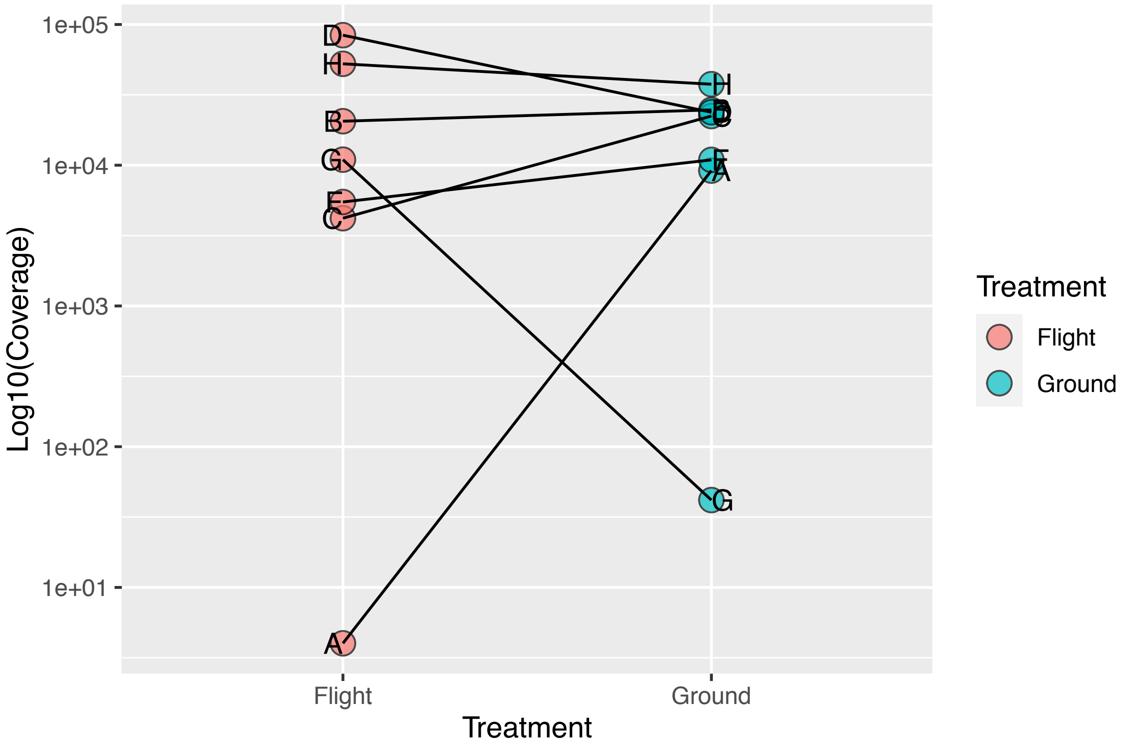

Figure **S-4**: Assessment of coverage for plasmid pSCR119 extracted from Flight and Ground samples, transformed into *E. coli*, then extracted and sequenced. All plasmid pairs with a mate had very high sequencing coverage.

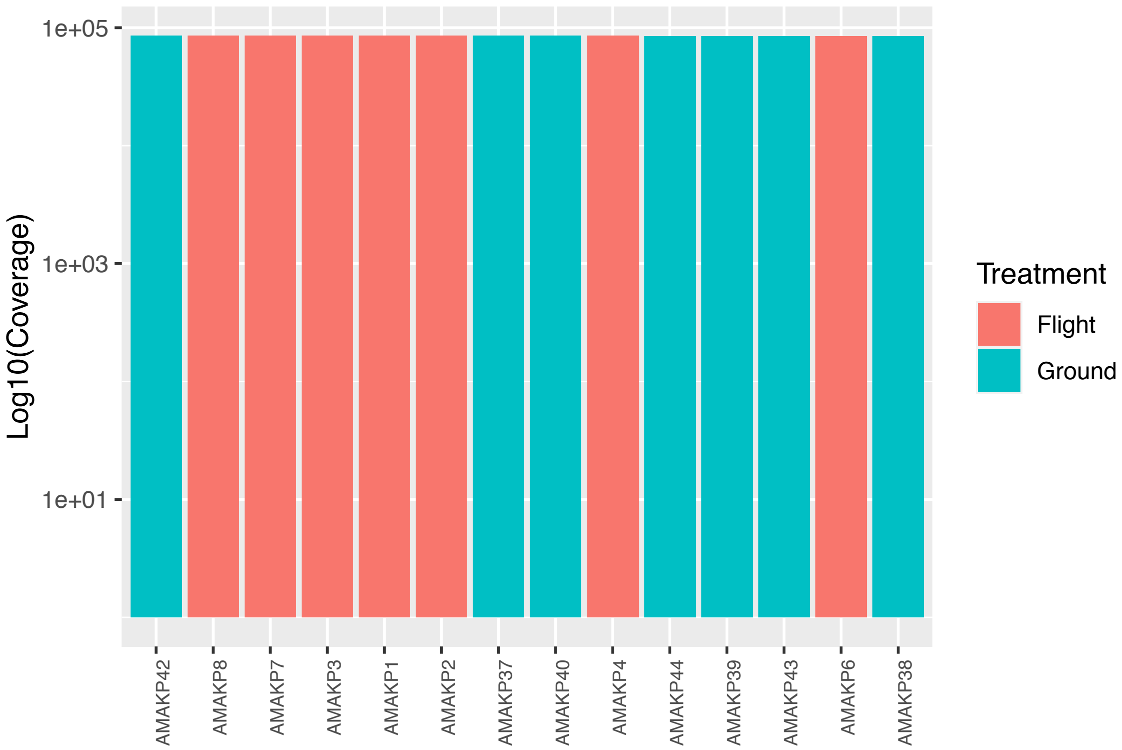

Figure **S-5**: Assessment of coverage for plasmid pSCR119 extracted from Flight and Ground samples, transformed into *E. coli*, then extracted and sequenced. All plasmid pairs with a mate had very high sequencing coverage and there was little difference in coverage among the sample pairs.

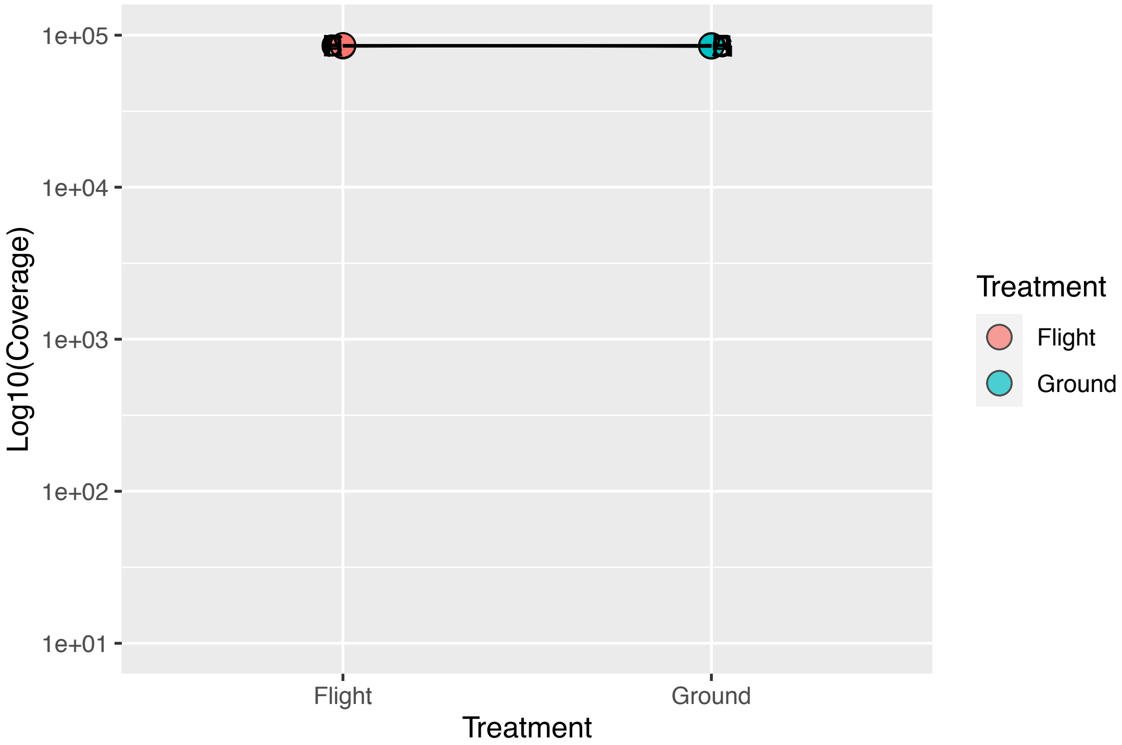

**Table S-1**: Growth media tested. For all the conditions in the table, 1 mL of *N. punctiforme* cryostock was thawed and added to 50 mL of BG11 media supplemented with different concentrations of $NaHCO_3$. Growth of the cultures was assessed at four time points by visual observation and Chl *a* extraction. Growth levels are ranked '-' to '+++' with '-' representing no growth detected and '+++' representing the highest growth detected.

| | Growth based on visual observation and Chl *a* extraction | | | |
|---|---|---|---|---|
| **$NaHCO_3$ concentration (mM)** | **Day 4** | **Day 6** | **Day 8** | **Day 10** |
| 0 | - | - | + | ++ |
| 1 | - | - | + | ++ |
| 2 | - | + | ++ | ++++ |
| 4 | - | + | ++ | +++ |
| 6 | - | - | ++ | ++ |
| 8 | - | - | + | ++ |
| 10 | - | - | - | + |

**Table S-2**: Transformation protocols and antibiotic concentration tested. Success rate labeled '-' represents unsuccessful transformations, '+' represent a success rate of up to 25%, '++' represents a success rate up to 75%, and '+++' represents a success rate up to 100%.

| Transformation method | Sonication | Plamid DNA added (µg) | [Antiobiotics] (µg/mL) | Success rate |
|---|---|---|---|---|
| Electroporation | 10 x 10 sec bursts | 5 | Kan 5 µg/mL | - |
| | 10 x 10 sec bursts | 10 | Kan 5 µg/mL | - |
| | 10 x 10 sec bursts | 5 | Kan 10 µg/mL | - |
| | 10 x 10 sec bursts | 10 | Kan 10 µg/mL | - |
| | 5 x 10 sec bursts | 5 | Kan 5 µg/mL | - |
| | 5 x 10 sec bursts | 10 | Kan 5 µg/mL | - |
| | 5 x 10 sec bursts | 5 | Kan 10 µg/mL | - |
| | 5 x 10 sec bursts | 10 | Kan 10 µg/mL | - |
| | 4 x 5 sec bursts | 5 | Kan 10 µg/mL | - |
| | 4 x 5 sec bursts | 5 | Kan 5 µg/mL | - |
| | 4 x 5 sec bursts | 5 | Kan 2.5 µg/mL | - |
| | 4 x 5 sec bursts | 5 | Kan 1.25 µg/mL | - |
| | 4 x 5 sec bursts | 10 | Kan 10 µg/mL | - |
| | 4 x 5 sec bursts | 10 | Kan 5 µg/mL | + |
| | 4 x 5 sec bursts | 10 | Kan 2.5 µg/mL | + |
| | 4 x 5 sec bursts | 10 | Kan 1.25 µg/mL | - |
| | 4 x 5 sec bursts | 15 | Kan 10 µg/mL | - |
| | 4 x 5 sec bursts | 15 | Kan 5 µg/mL | - |
| | 4 x 5 sec bursts | 15 | Kan 2.5 µg/mL | - |
| | 4 x 5 sec bursts | 15 | Kan 1.25 µg/mL | - |
| | 4 x 5 sec bursts | 5 | Neo 10 µg/mL | - |
| | 4 x 5 sec bursts | 5 | Neo 5 µg/mL | - |
| | 4 x 5 sec bursts | 5 | Neo 2.5 µg/mL | - |
| | 4 x 5 sec bursts | 5 | Neo 1.25 µg/mL | - |
| | 4 x 5 sec bursts | 10 | Neo 10 µg/mL | - |
| | 4 x 5 sec bursts | 10 | Neo 5 µg/mL | + |
| | 4 x 5 sec bursts | 10 | Neo 2.5 µg/mL | + |
| | 4 x 5 sec bursts | 10 | Neo 1.25 µg/mL | - |
| | 4 x 5 sec bursts | 15 | Neo 10 µg/mL | - |
| | 4 x 5 sec bursts | 15 | Neo 5 µg/mL | - |
| | 4 x 5 sec bursts | 15 | Neo 2.5 µg/mL | - |
| | 4 x 5 sec bursts | 15 | Neo 1.25 µg/mL | - |
| | 3 x 5 sec bursts | 5 | Kan 10 µg/mL | - |
| | 3 x 5 sec bursts | 5 | Kan 5 µg/mL | ++ |
| | 3 x 5 sec bursts | 5 | Kan 2.5 µg/mL | ++ |
| | 3 x 5 sec bursts | 5 | Kan 1.25 µg/mL | + |
| | 3 x 5 sec bursts | 10 | Kan 10 µg/mL | - |
| | 3 x 5 sec bursts | 10 | Kan 5 µg/mL | +++ |
| | 3 x 5 sec bursts | 10 | Kan 2.5 µg/mL | +++ |
| | 3 x 5 sec bursts | 10 | Kan 1.25 µg/mL | ++ |
| | 3 x 5 sec bursts | 5 | Neo 10 µg/mL | - |
| | 3 x 5 sec bursts | 5 | Neo 5 µg/mL | ++ |
| | 3 x 5 sec bursts | 5 | Neo 2.5 µg/mL | ++ |
| | 3 x 5 sec bursts | 5 | Neo 1.25 µg/mL | + |
| | 3 x 5 sec bursts | 10 | Neo 10 µg/mL | - |
| | 3 x 5 sec bursts | 10 | Neo 5 µg/mL | +++ |
| | 3 x 5 sec bursts | 10 | Neo 2.5 µg/mL | +++ |
| | 3 x 5 sec bursts | 10 | Neo 1.25 µg/mL | ++ |

**Table S-3**: Number of colonies obtained after transforming raw DNA extracted from Ground and Flight samples into *E. coli* as described in the Methods section.

| Sample | Number of colonies per plate |
|---|---|
| 1 | 26 |
| 2 | 1076 |
| 3 | 444 |
| 4 | 77 |
| 5 | 0 |
| 6 | 524 |
| 7 | 79 |
| 8 | 28 |
| 37 | 5 |
| 38 | 1720 |
| 39 | 320 |
| 40 | 60 |
| 41 | 0 |
| 42 | 1700 |
| 43 | 636 |
| 44 | 1040 |

**Table S-4**: DNA concentration sent for sequencing. Sample numbering is the same as shown in Table 1.

| Total DNA | Sample label | [raw DNA] (ng/µL) | Sample volume (µL) | DNA (ng) | 260/280 | 260/230 |
|---|---|---|---|---|---|---|
| Flight | 1 | 33.7 | 35 | 1179.5 | 1.84 | 2.41 |
| | 2 | 46.0 | 35 | 1610.0 | 1.9 | 2.4 |
| | 3 | 13.1 | 35 | 458.5 | 1.84 | 2.4 |
| | 4 | 11.5 | 35 | 402.5 | 1.88 | 2.45 |
| | 5 | 5.8 | 35 | 202.0 | 1.9 | 2.4 |
| | 6 | 13.9 | 35 | 486.5 | 1.86 | 2.36 |
| | 7 | 9.9 | 35 | 345.5 | 1.9 | 2.4 |
| | 8 | 32.0 | 35 | 1120.0 | 1.9 | 2.4 |
| Ground | 37 | 11.6 | 35 | 406.0 | 1.9 | 2.4 |
| | 38 | 45.5 | 35 | 1592.5 | 1.84 | 2.35 |
| | 39 | 6.1 | 35 | 214.9 | 1.85 | 2.39 |
| | 40 | 8.5 | 35 | 296.5 | 1.9 | 2.4 |
| | 41 | 5.1 | 35 | 178.5 | 1.85 | 2.42 |
| | 42 | 7.9 | 35 | 276.9 | 1.9 | 2.4 |
| | 43 | 35.3 | 35 | 1235.5 | 1.9 | 2.4 |
| | 44 | 38.8 | 35 | 1358.0 | 1.8 | 2.4 |

| Plasmid DNA | Sample label | [raw DNA] (ng/µL) | Sample volume (µL) | DNA (ng) | 260/280 | 260/230 |
|---|---|---|---|---|---|---|
| Flight | P1 | 50.0 | 45 | 2250.0 | 1.86 | 2.32 |
| | P2 | 359.0 | 45 | 16155.0 | 1.88 | 2.34 |
| | P3 | 264.0 | 45 | 11880.0 | 1.88 | 2.36 |
| | P4 | 319.0 | 45 | 14355.0 | 1.86 | 2.26 |
| | P6 | 200.0 | 45 | 9000.0 | 1.89 | 2.59 |
| | P7 | 81.0 | 45 | 3645.0 | 1.93 | 2.48 |
| | P8 | 85.7 | 45 | 3856.5 | 1.87 | 2.33 |
| Ground | P37 | 216.0 | 45 | 9720.0 | 1.86 | 233 |
| | P38 | 334.0 | 45 | 15030.0 | 1.86 | 2.38 |
| | P39 | 127.0 | 45 | 5715.0 | 1.84 | 2.28 |
| | P40 | 192.0 | 45 | 8640.0 | 1.87 | 2.35 |
| | P42 | 364.0 | 45 | 16380.0 | 1.88 | 2.38 |
| | P43 | 238.0 | 45 | 10710.0 | 1.88 | 2.38 |
| | P44 | 330.0 | 45 | 14850.0 | 1.89 | 2.37 |
| pSCR119 positive control from transformation | P119 | 240.0 | 45 | 10800.0 | 1.85 | 2.35 |

**Supplementary Information S-1**: R script for outputs from inStrain analysis will be provided upon request.
